# Revisiting the expression signature of *pks15/1* unveils regulatory patterns controlling phenolphtiocerol and phenolglycolipid production in pathogenic mycobacteria

**DOI:** 10.1101/2020.02.20.950329

**Authors:** Beatriz Ramos, Stephen V. Gordon, Mónica V. Cunha

**Author notes:** Correspondence: Mónica V. Cunha - National Institute for Agrarian and Veterinary Research (INIAV, IP), Oeiras, Portugal.

## Abstract

One of the most relevant and exclusive characteristics of mycobacteria is its cell wall, composed by mycolic acids. Amid these are two related families of glycosylated lipids, diphthioceranates and phthiocerol dimycocerosate (PDIM) and its variant phenolic glycolipids (PGL). PGL have been associated with cell wall impermeability, phagocytosis, defence against nitrosative and oxidative stress and, supposedly, biofilm formation. In bacteria from the *Mycobacterium tuberculosis* complex, the biosynthetic pathway of the phenolphthiocerol moiety of PGL depends upon the expression of several genes encoding type I polyketide synthases (PKS), namely *ppsA-E* and *pks15/1* constituting the PDIM + PGL locus, highly conserved in PDIM/PGL-producing strains. Consensus has not been achieved regarding the genetic organization of *pks15/1* locus and little effort has been put on the disclosure of its transcriptional signature. Here we explore publicly available datasets of transcriptome data (RNA-seq) from more than 100 experiments in 40 growth conditions to outline the transcriptional structure and signature of *pks15/1* and use a differential expression approach to infer the regulatory patterns involving these and related genes. We show that *pks1* is highly correlated with *fadD22, Rv2949c, lppX, fadD29* and, also, *pks6* and *pks12*, with the first three putatively integrating a polycistronic structure. We evidence dynamic heterogeneity of transcription within the genes involved in phenolphtiocerol and phenolglycolipid production, most exhibiting up-regulation upon acidic pH and antibiotic exposure and down-regulation under hypoxia, dormancy, and low/high iron concentration. We finally propose a model based on transcriptome data in which σ^D^ positively regulates *pks1, pks15* and *fadD22*, while σ^B^ and σ^E^ factors exert negative regulation at an upper level.

## 1 Introduction

Tuberculosis (TB) is an infectious disease caused by *Mycobacterium tuberculosis* (*Mtb)* that remains a major public health concern. In 2016, approximately 6.3 million new cases of TB were reported (1). The mycobacterial cell wall is a crucial interface of *Mtb* and other pathogenic mycobacteria with the host wherein *Mycobacterium*-specific components are located (2). Recently, Chiaradia and coworkers (2017) proposed a cell wall structure composed of three layers, namely the mycomembrane, arabinoglactan and peptidoglycan. This model proposes that the inner leaflet of the mycomembrane is composed by mycolic acids that are esterified to arabinogalactan, which in turn is covalently attached to peptidoglycan (3). Amid the *Mycobacterium-*specific components are two related families of glycosylated lipids: diphthioceranates (DIP) and phthiocerol dimycocerosate (PDIM), along with its variant phenolic glycolipids (PGL) (2). PGL are known to be associated with several cellular functions, namely impermeability of the cell wall, phagocytosis (4, 5), defence mechanisms against nitrosative and oxidative stress (6) and, theoretically, to the ability of mycobacteria to form biofilms (7).

In this work, we focus on the transcription signature of genes comprised in the biosynthetic pathway responsible for the synthesis of the phenolphthiocerol moiety of PGL when *Mtb* is grown under a different set of stressors mimicking the host environment, namely pH variation, different carbon sources, limiting or excessive iron concentration, hypoxia, dormancy, phosphate depletion and antibiotic exposure. The genes related to the biosynthesis of PGL belong to the class of polyketide synthases (PKS). There are three types of PKS, classified according to their structure and biosynthetic function. Type I PKS contain multiple catalytic domains and can be classified as modular or iterative. Modular type I PKS have distinct functional domains that are used only once during the formation of the product. On the other hand, iterative PKS have functional domains that intervene repetitively to produce the final polyketide. Type II PKS are composed by several enzymes, each one carrying a single and distinct catalytic domain that is used iteratively during formation of the polyketide product. Chalcone synthase-like PKS, the type III PKS, represent a more divergent group that, in opposition to types I and II PKS, do not require the involvement of acyl carrier proteins (ACP) (8). Among the genes required for PGL production, are *ppsA-E* and *pks15/1* encoding type I PKS, constituting the PDIM + PGL *locus*, which is known to be highly conserved in PDIM/PGL-producing strains (9). Type I PKS modules are constituted by a minimal set of three domains, namely a ketoacyl synthase (KS) domain, an acyltransferase (AT) domain, and an acylcarrier protein (ACP) domain. This module can also contain one or more of the following domains: keto reductase (KR), dehydratase (DH) and enoyl reductase (ER) (10). The *pks15* encodes a KS domain while *pks1* encodes KR, DH, ER, AT and ACP domains. It is reported that a 7 bp deletion in some *Mtb* strains and a 1 bp deletion in a few *Mb* strains leads to a frameshift that results in the split of *pks15* and *pks1* (11). Constant and co-workers (2002) documented that, in PGL-producers, the *pks1* and *pks15* correspond to a single gene, *pks15/1*, while they are apart (2 ORFs) in PGL-deficient strains like H37Rv or Erdman. PGL’s phenolphthiocerol moiety production starts with the enzyme encoded by *Rv2949c* that catalyses the formation of p-hydroxybenzoic acid (p-HBA) that will later be activated by *fadD22* product, with p-hydroxybenzoyl-AMP ligase activity, and finally elongated with malonyl-CoA as extender unit by *pks15/1*, in a reaction that may comprise eight to nine elongation cycles. The product of *fadD29*, a fatty acyl-AMP ligase, is then responsible for activation of p-hydroxyphenylalkanoates, later transferred onto the *ppsA* product and finally elongated with malonyl-CoA and mehtylmalonyl-CoA by PpsB-PpsE to yield the phenolphtiocerol moiety of PGL (12-14).

As described (15), upon the entrance of *Mtb* in its target cells, a cascade of events is promoted by the immune system that, in an immunocompetent host, leads to granuloma formation and bacterial enclosure. This structure is beneficial for the host, since it confines infection to localized regions, preventing bacterial spreading. Several conditions like starvation, presence of reactive oxygen and nitrogen intermediates, or hypoxia inside granulomas, limited amount of iron, the inorganic phosphate (P_i_) scarcity and the typical low pH are some of the stresses mycobacteria are exposed to during infection (15).

Along with stress-induced genes, also described in *Mtb* is a group of 13 σ subunits responsible for transcriptional regulation, namely the essential housekeeping sigma factor (σ^A^), the stress-responsive factor (σ^B^) and the other 11 sigma factors described so far that act as environmental responsive regulators (σ^C-M^). Several studies have been performed to infer the role of each sigma factor and the condition that triggers their activation, initially by analysing expression levels and, more recently, by the construction of deletion strains (16, 17). The presence and articulation of these wide variety of sigma factors enables an adaptive transcriptional response for a large set of conditions. Chauhan and co-workers (2016) performed a reconstruction of the sigma factor regulatory network that enabled the clarification of the direct and indirect connections among the 13 factors (18). This study has defined an hierarchical organization of sigma factors in *Mtb*, as well as the definition of multiple factor usage to respond to specific stresses. Current knowledge advocates a hierarchical organization that comprises three regulation levels: (i) top level: *sigA, sigB, sigH, sigM*; (ii) middle level: *sigE, sigF, sigG, sigJ, sigL*; and (iii) bottom level: *sigC, sigD, sigI, sigK*.

Besides host-induced stress, *Mtb* is frequently exposed to drug-induced stress as part of antibiotic therapy. For treatment of drug-susceptible TB, a combination of isoniazid, ethambutol, rifampicin, streptomycin or pyrazinamide is foreseen (19). Since TB treatment is a long-lasting process, *Mtb’s* transcriptional profile is expected to suffer severe changes along this process. *In vitro* studies have shown that for each of the above mentioned drugs, combined expression of a set of genes results in the antibiotic resistance phenotype of *Mtb*.

Due to their impact in the modulation of *Mtb* and *Mb* cell surfaces, which act as the interface with the host cell and exert a potential effect on pathogenicity, understanding the transcriptional profiles and transcriptional structure of *pks15’* and *pks1* under different stress conditions that mimic the host environment is of major importance. As such, this work aimed to pinpoint the regulatory patterns responsible for controlling *pks1* and *pks15* transcription by exploring publicly available large datasets of transcriptome data (RNA-seq). This methodological approach for transcriptome profiling takes advantage of deep-sequencing technologies to get a precise transcript measurement at the whole genome level. Our approach enabled us to define sets of correlated genes according to their expression profiles under different stress conditions by means of a differential expression approach, and also to outline the transcriptional structure of the *pks15/1* locus based on the available experimental data and *in silico* predictions.

## 2 Methods

### 2.1 *In silico* analysis of regulatory data of *pks1* and *pks15*

Regulatory information on *pks1* and *pks15* were gathered from international databases such as: Mycobrowser (20), National Center for Biotechnology Information (NCBI) (21), TB Database (22, 23) and MTB Network Portal (24) (visited from 09/2018 to 01/2019). MTB Network Portal reports information produced by the cMonkey algorithm that demonstrates that *pks1* and *pks15* belong to the same two biclusters. Biclusters are sets of co-regulated genes defined by cMonkey according to mRNA-based expression levels, *de novo* identification of transcription factor binding motifs and pre-established association pathways. Location of putative ribosomal binding sites (RBS) was inferred for *pks1* with *Prokaryotic Dynamic Programming Gene finding Algorithm* (PRODIGAL) (25). Synteny analyses were performed for *pks1, pks15* and *fadD22* using SyntTax (Prokaryotic Synteny & Taxonomy Explorer) (26).

### 2.2 RNA-seq data and differential expression analyses of a selected panel of genes

For expression analyses, 105 experiments from *Mtb* strains (*Mtb* H37Rv and *Mtb* CDC1551), constituting a set of 40 experimental conditions in seven datasets (Accession codes at NCBI: GSE47863, GSE67035, GSE52020, GSE83814, GSE66408, GSE104599 and GSE107831) were considered. This analysis was performed for a set of 90 genes, including *pks1, pks15, fadD22, fadD29* genes comprised in bicluster modules 0211 and 0490 from *MTB* Network Portal that represent co-regulated genes, genes encoding PKS and σ factors, and genes encoding regulatory factors for each of the experimental conditions. Regarding *M. bovis* BCG, 21 experiments from *M. bovis* BCG str. Pasteur 1173P2 constituting a set of seven experimental conditions in two datasets (accession codes at NCBI: GSE66883 and GSM3160698) were analysed. For this analysis, a set of 50 genes were selected, including *pks1, pks15, fadD22, fadD29* genes comprised in bicluster modules 0211 and 0490 from *MTB* Network Portal, that represent co-regulated genes, and genes encoding PKS and σ factors. For each experiment, reads were extracted in FASTQ format using NCBI SRA Toolkit v.2.8.1.3 (27). Those FASTQ files were mapped against a reference genome, *Mtb* H37Rv (RefSeq code: NC_000962.3, version 3) with TopHat v.2.1.0.54, (28, 29), using default settings to produce a BAM file containing a list of read alignments. Transcript identification and counting was later performed with *bias* correction by Cufflinks v.2.2.1.0 (28, 30) using as reference the annotation of the genomes listed above. Cufflinks was used to calculate the relative abundance of each gene in *Reads Per Kilobase per Million mapped reads* (RPKM). The RPKM values were transformed by log_10_, heatmaps were plotted by GraphPad Prism 7(31) and dendrograms were computed using BioNumerics v4.0 (Applied Maths) calculating Pearson correlation coefficient and the *unweighted pair group method with arithmetic means* (UPGMA) as the agglomerative clustering algorithm. Pearson correlation coefficient was calculated using GraphPad Prism 7 and correlation network was plotted using Cytoscape v.3.5.1(32, 33). For differential expression analysis, htseq-count v.0.9.1(34) was used to count reads mapped to each gene and DESeq v.2.11.40.1 (35) was used to determine differentially expressed genes from count tables using Wald statistic test with *p*-value adjusted for multiple testing with the Benjamini-Hochberg procedure (α=0.05). For evaluation of significance it was considered the following scale: significant (*p*-value=0.01 to 0.05); very significant (*p*-value=0.001 to 0.01); extremely significant (*p*-value=0.0001 to 0.001); extremely significant (*p*-value< 0.0001). Data was plotted as heatmap using GraphPad Prism 7. The public server at usegalaxy.org (36) was used to analyse the data with NCBI SRA Toolkit, TopHat, Cufflinks, htseq-count and DESeq2.

## 3 Results and Discussion

### 3.1 Revisiting the organization of *pks1* and *pks15* genetic locus across *Mtb* genomes based on predicted regulatory data and homology searches

Encoded in the minus strand, from position 3291503 to 3296353 for *pks1* and, from position 3296350 to 3297840 for *pks15*, the *Mtb pks1* and *pks15* genes together encode a polyketide synthase with six identified domains involved in the synthesis of PGL. They appear to have a functional cooperation with *fadD22*, a bidomain initiation module. In the MTB Network Portal (retrieved on January, 2019), the *pks1* and *pks15* genes are placed together in bicluster modules 0211 and 0490, with residual values of 0.5 and 0.57, respectively, meaning that bicluster module 0211 presents a tighter expression profile amongst its members, which should also indicate better bicluster quality and thus more certainty associated with co-expression. Also related are the two genes upstream of *pks15, fadD22*, included in the same two modules, and *Rv2949c*, included in module 0490 (Figure 1). The mRNA-based expression levels, *de novo* identification of transcription factor binding motifs and pre-established association pathways, each support that both *pks1* and *pks15* may be regulated by the products of seven genes: positively by *Rv0042c, sigK, Rv2258c* and *Rv3557c*; negatively by *sigB, Rv2745c* and *Rv3583c*. Furthermore, according to chIP-seq data, *pks1* is bound by the transcription factor *Rv3830c* with no differential expression reported. The operon structure is undetermined: TB Database suggests four different combinations of six genes (*fadD29, Rv2949c, fadD22, pks15, pks1* and *lppX*), while MTB Network Portal suggests an operon composed by five genes: *fadD29, Rv2949c, fadD22, pks15* and *pks1*. All those genes are involved in the biosynthesis of phenolphthiocerol moiety of PGL, except *lppX* that was shown to be involved in the translocation of PDIM to the outer membrane (37).

**Figure 1.**
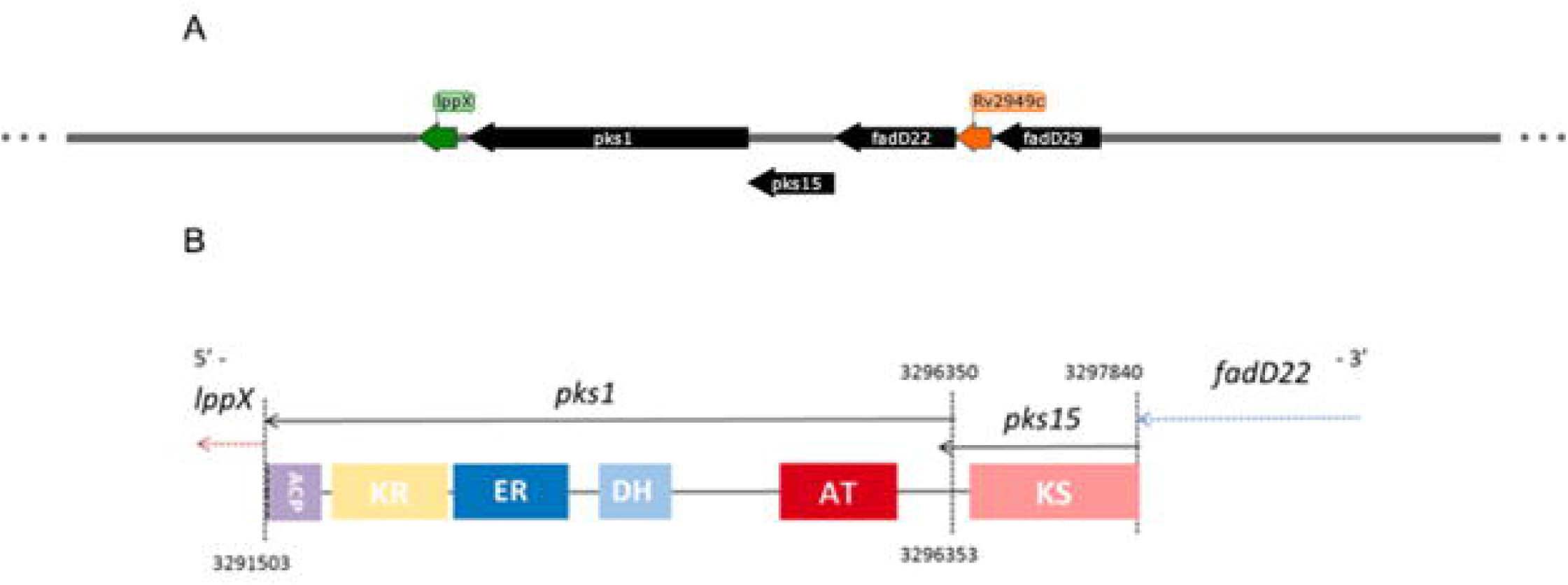
Genomic locus of *pks1* and *pks15*, protein domains and their role in the biosynthetic pathway of PGL. A – Schematic representation of the location of *pks1* and *pks15* in the minus strand of *M. tuberculosis* H37Rv genome. In black: lipid metabolism. In green: cell wall and cell processes. In orange: intermediary metabolism and respiration. B – Domain organization of Pks1 and Pks15. Abbreviations: KS, ketoacylsynthase; AT, acyltransferase; DH, dehydratase; ER, enoylreductase; KR, ketoreductase; ACP, acylcarrier protein.

To compare the conservation of *pks15/1* locus across *Mtb* genomes, a synteny analysis was performed using the Pks1 sequence from *Mtb* H37Rv as the query. Among the 210 *Mtb* accession codes available, those with synteny scores above 80 were 90.5% and 99.5% for *pks15* and *fadD22*, respectively. For *pks1*, 89.5% of the accession codes used for analysis presented a score higher than 80 (Figure 2). When analysing local genomic conservation, it was possible to denote an irregular pattern across Pks1 and Pks15/1, while Rv2949c, FadD22, LppX, Rv2944 and Rv2943 presented a regular organization pattern across most genomes analysed. However, when comparing the top three scored genomes, it was also possible to identify that, for *Mtb* 0B070XDR (GenBank code: CP008970.1), the element upstream of Rv2949c does not present homology with FadD29, whilst homology was found for a protein encoded on the opposite strand of *pks1*.

**Figure 2.**
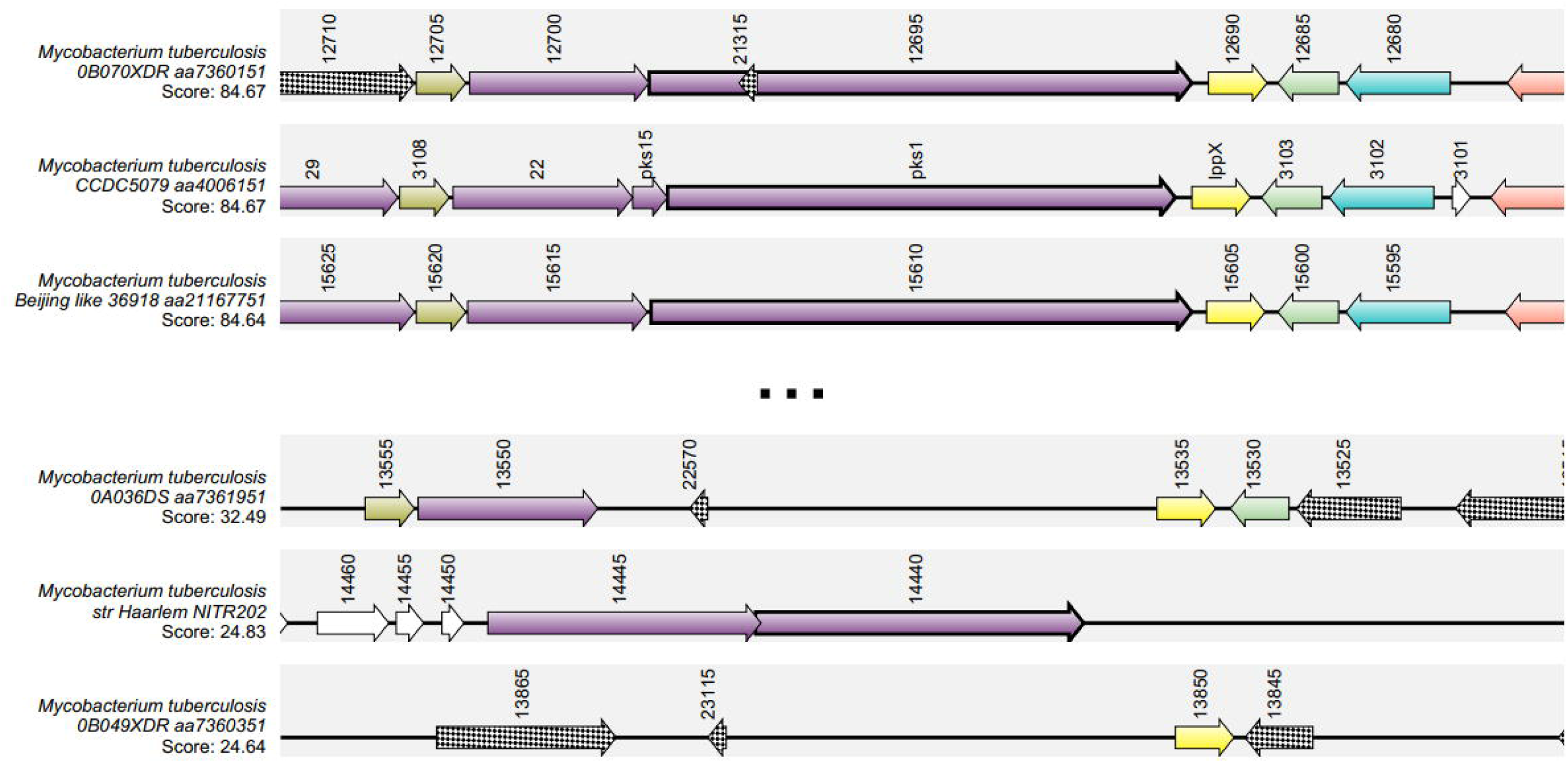
Representation of top three and bottom three scores from synteny analysis. Top 3 and bottom 3 synteny scores for *pks1* as predicted by SyntTax. Color code represents matching proteins across genomes.

### 3.2 Analysis of expression data for a selected panel of genes enclosing *pks1* and *pks15*

To characterize the transcriptional signature of *pks1* and of presumably correlated genes, RNA-seq based expression analyses focused on a set of 90 genes, including *pks1, pks15, fadD22, fadD29*, other genes encoding polyketide synthases and σ factors, were scrutinized. Transcriptional profiles were compared across a set of 40 experimental conditions. Data gathered was analysed by alignment against a reference genome [*Mtb* H37Rv (RefSeq code: NC_000962.3, version 3)], by read counting and through the calculation of RPKM as a proxy for gene expression in each condition. RPKM is a relative value, meaning that it varies according not only to read count, but also to the total number of reads obtained for each experiment, allowing comparison across experiments.

To understand the similarity across RPKM profiles, a dendrogram was generated, using 85% similarity as a cut-off for cluster formation. This cut-off was established to include the top quarter of similarity values in this analysis. We then obtained a total of 47 clusters, 36 being single member clusters. Cluster VI includes some of the genes of interest (*lppX, pks1, fadD22* and *Rv2949c*) along with *pks6*, which is a *pks1* paralog (Figure 3). Another relevant cluster (IX) includes *papA1, papA3, pks2* and *pks4*, a group of genes known to be upregulated at low pH (Figure 3). The larger cluster (III) includes most genes responsible for antibiotic resistance and regulatory factors, along with *fadD29* (Figure 3). To obtain a more focused approach on direct interactions between genes, we constructed a correlation network with pairs of genes exhibiting correlation factors above 0.85. In this network, it becomes evident that *pks1* is highly correlated with *lppX* (Pearson correlation coefficient of 0.881), *fadD22* (0.902), *Rv2949c* (0.885), *fadD29* (0.859) and also with *pks6* (0.895) and *pks12* (0.880) (Figure 4A). Unexpectedly, some RPKM values registered for *pks15* were null, possibly due to the absence of mapped reads, and thus were not plotted in this network.

**Figure 3.**
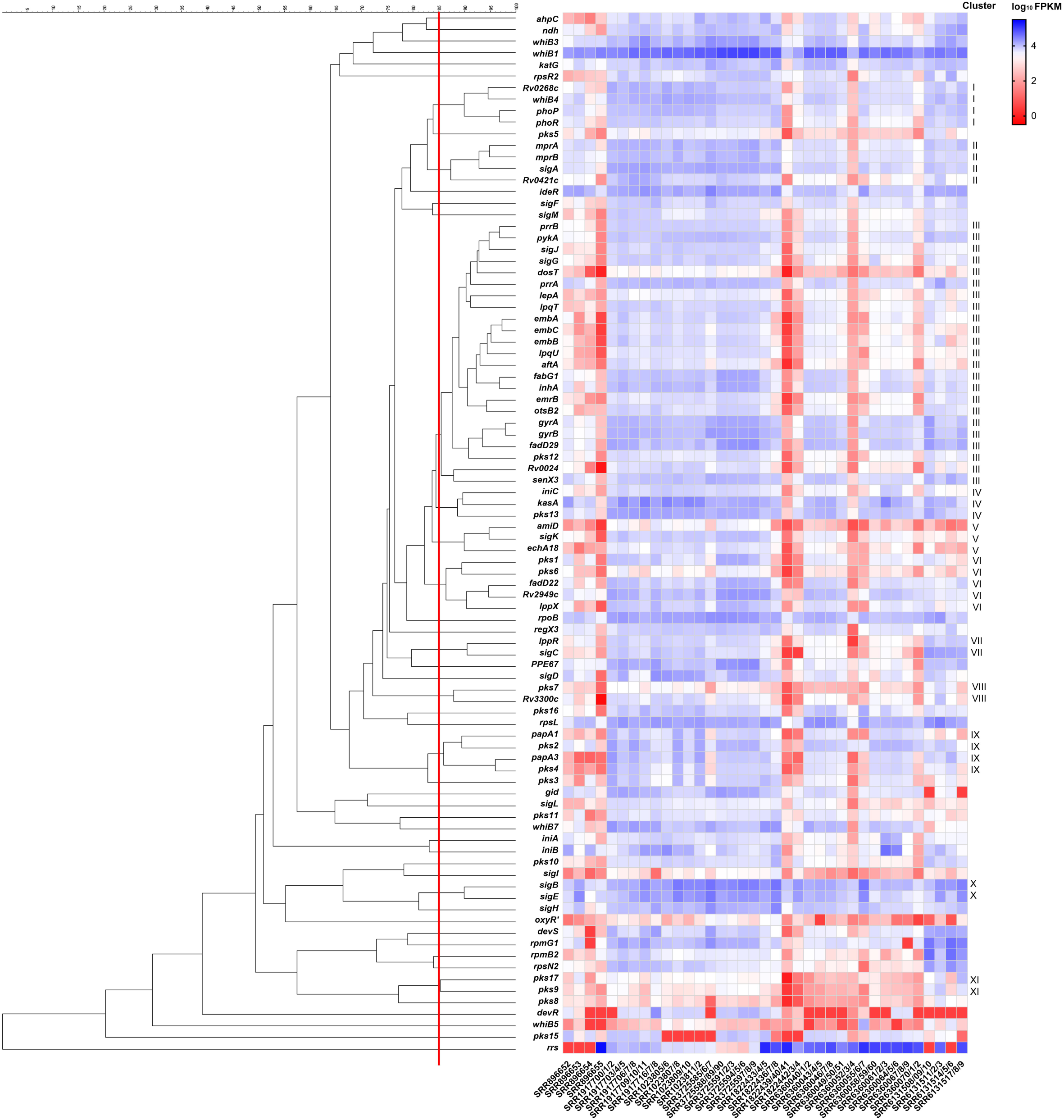
Expression profiling of selected genes from *Mycobacterium tuberculosis*, presented as log_10_ RPKM. Cut-off: 85% of similarity. Abbreviations: SRR896652 - *Mtb* H37Rv grown in dextrose at exponential phase; SRR896653 – *Mtb* H37Rv grown in dextrose at stationary phase; SRR896654 - *Mtb* H37Rv grown in long fatty acids at exponential phase; SRR896655 - *Mtb* H37Rv grown in long fatty acids at stationary phase; SRR1917700/1/2 - *Mtb* H37Rv grown in 0.4% glucose; SRR1917703/4/5 - *Mtb* H37Rv grown in high iron concentration; SRR1917706/7/8 - *Mtb* H37Rv grown in low iron concentration for 1 day; SRR1917709/10/11 - *Mtb* H37Rv grown in low iron concentration for 1 week; SRR1917716/7/8 - *Mtb* HN878; SRR1023805/6 - *Mtb* CDC1551 grown in glycerol at pH 7; SRR1023807/8 - *Mtb* CDC1551 grown in glycerol at pH 5.7; SRR1023809/10 - *Mtb* CDC1551 grown in pyruvate at pH 7; SRR1023811/2 - *Mtb* CDC1551 grown in pyruvate at pH 5.7; SRR3725585/6/7 - *Mtb* H37Rv grown in hypoxia; SRR3725588/89/90, SRR3725591/2/3, SRR3725594/5/6, SRR3725597/8/9 - *Mtb* H37Rv (1-4) day(s) after reaeration; SRR1822433/4/5 - *Mtb* H37Rv in log phase; SRR1822436/7/8 - *Mtb* H37Rv in early dormancy phase; SRR1822439/40/41 – *Mtb* H37Rv in medium dormancy phase; SRR1822442/3/4 - *Mtb* H37Rv in late dormancy phase; SRR6360040/1/2 – *Mtb* H37Rv grown with CIP for 4h; SRR6360043/4/5 - *Mtb* H37Rv grown with INH for 4h; SRR6360046/7/8 - *Mtb* H37Rv grown with EMB for 4h; SRR6360049/50/51 - *Mtb* H37Rv grown with STR for 4h; SRR6360052/3/4 - *Mtb* H37Rv grown with RIF for 4h; SRR6360055/6/7 - *Mtb* H37Rv grown in control conditions; SRR6360058/59/60 - *Mtb* H37Rv grown with CIP for 24h; SRR6360061/2/3 - *Mtb* H37Rv grown with INH for 24h; SRR6360064/5/6 - *Mtb* H37Rv grown with EMB for 24h; SRR6360067/8/9 - *Mtb* H37Rv grown with STR for 24h; ; SRR6360070/1/2 - *Mtb* H37Rv grown with RIF for 24h; SRR6131508/09/10 - *Mtb* H37Rv grown in control conditions; SRR6131511/2/3 - *Mtb* H37Rv grown with PBS; SRR6131514/5/6 - *Mtb* H37Rv grown in phosphate depletion; and SRR6131517/8/9 - *Mtb* H37Rv grown in hypoxia.

**Figure 4.**
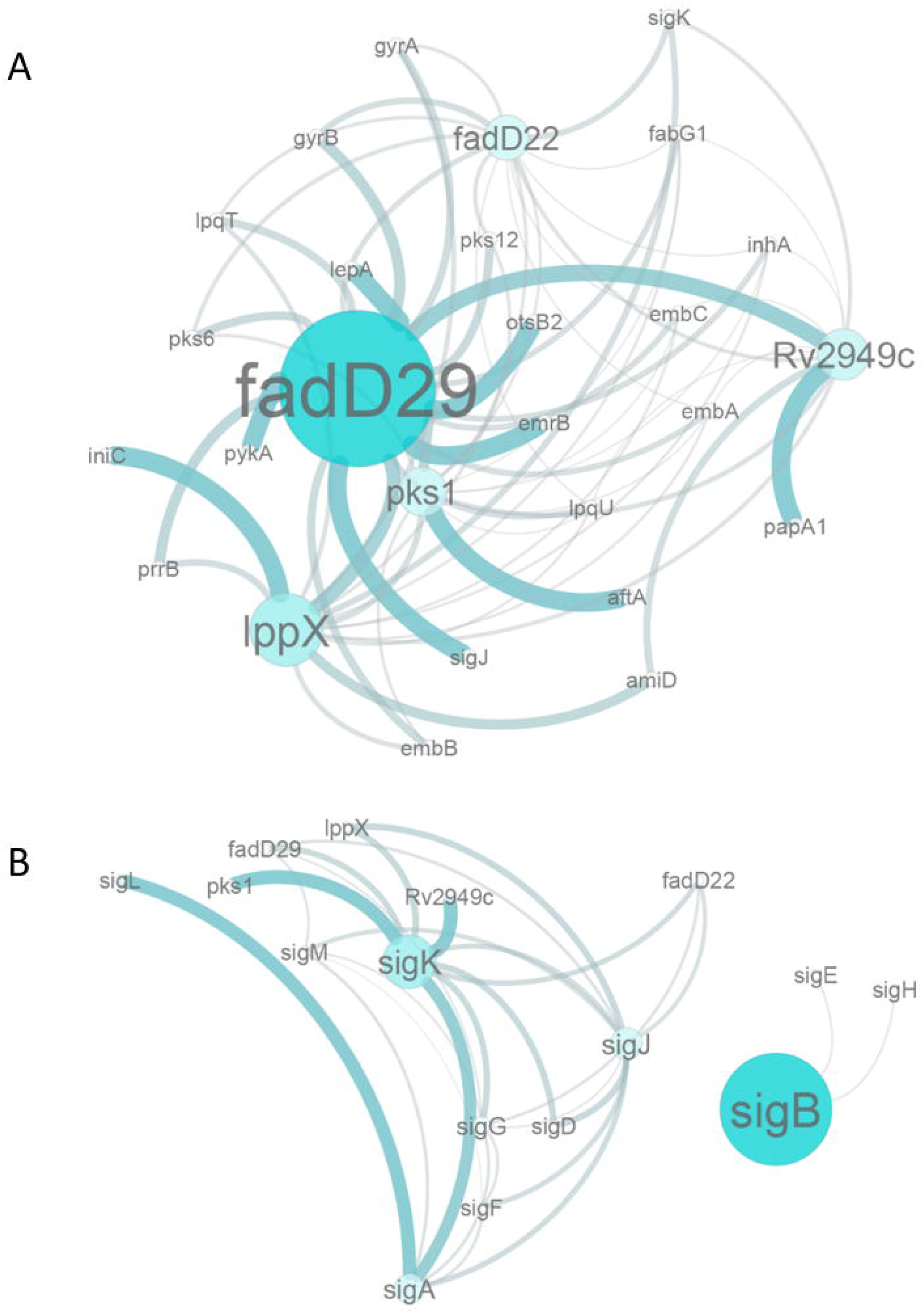
Correlation network of expression data. A: Correlation threshold=0.85. B: Correlation threshold=0.8. Thicker connections represent stronger correlations.

A correlation between σ factors could also be confirmed. The *sigA* gene, known as the housekeeping regulator, is correlated with *sigG* (0.868), *sigJ* (0.902) and *sigK* (0.871); *sigB* is correlated with *sigE* (0.900); *sigG* with *sigJ* (0.926), *sigK* (0.879) and *sigM* (0.852); and *sigJ* with *sigK* (0.920) (Figure 4A). Both analyses provide indication that *pks1* expression is highly correlated with the expression profiles of *fadD22, Rv2949c* and *fadD29*, in agreement with reports from microarray data (38). The expression pattern of *pks15* revealed by this analysis is impressively different from the one found in *pks1* in some of the stress conditions under examination, turning *pks15* into a single member cluster placed apart from the remaining clusters and, consequently, being absent from the correlation network. In contrast with these results, microarray data available online (49) suggest that *pks15* is also highly correlated with *pks1* and *fadD22*. Our analyses also suggest that *pks4* is correlated with *pks3*, once their expression profiles share 82.9% similarity (Figure 3), thus agreeing with previously published data (39) reporting that a *Mtb* H37Rv double mutant for *pks4* and *pks3* is not able to produce mycolipanoic, mycolipenic, and mycolipodienoic acids. Also, the fact that *pks3* and *pks4* form a polyketide structure similar to *pks15* and *pks1*, respectively, wherein *pks3* and *pks15* both encode the ketoacyl synthase domain and *pks4* and *pks1* both encode the remaining polyketide synthase domains, would suggest that *pks15* and *pks1* could also be highly correlated. Naturally, we were expecting to confirm this correlation across the selected experimental datasets.

Even though gene expression does not necessarily represent the activity of a specific σ factor, we integrated this correlation network with sigma factor expression data to plot a representation of the putative regulation of selected genes by σ factors. Eleven of the 13 sigma genes under analysis are highly correlated. Both *sigA*, the housekeeping σ factor of mycobacteria, and *sigM*, responsible for long-term adaptation, are known to be regulated by *sigC*, whose role is to control virulence and immunopathology. The *sigA* factor is also known to regulate *sigG*, mostly induced during macrophage infection, which will regulate *sigJ*, known to be overexpressed in late stationary phase of dormant cultures. The *sigG* and *sigJ* will further regulate *sigL*, known to be involved in PDIM biosynthesis, that in turn regulate *sigK*, whose role remains partially unclear (17). Some correlations with sigma factors plotted here directly involve the selected panel of genes, namely between *sigJ* and *fadD29* (0.851) and *sigK* with *lppX* (0.891), *fadD22* (0.852) and *Rv2949c* (0.879) (Figure 4B). The *sigK* factor is predicted by *in silico* analysis to positively regulate *pks1* and *pks15* (40). At a lower correlation threshold, this predicted regulation becomes clearer, as *sigK* presents correlation with *pks1* (0.835) (data not shown). On the contrary, amongst the analyses focused on sigma factors, *sigE* was the factor that presented the lowest correlations with the established genes of interest (*lppX*, 0.242; *pks1*, 0.010; *fadD22*, 0.202; *Rv2949c*, 0.205; and *fadD29*, 0.261) (data not shown).

For *M. bovis* BCG, we obtained a total of 14 clusters, with five being single member clusters using 80% similarity as cut-off. Among those, it was possible to identify a cluster comprising *Mb2973c, pks13, fadD22, pks15/1, lppX, sigMa* and *sigC*, sharing 86% similarity (Figure 5). In *M. bovis* BCG, as verified for *M. tuberculosis, pks15/1* and *fadD22* exhibit a correlation value of 0.969, with correlation values between *pks15/1* and *fadD22* and *lppX* and *Mb2973c* all above 0.9. In addition, *lppX, pks15/1, fadD22* and *Mb2973c* were also shown to be correlated with *sigA* (above 0.9) and with *sigC* (above 0.869).

**Figure 5.**
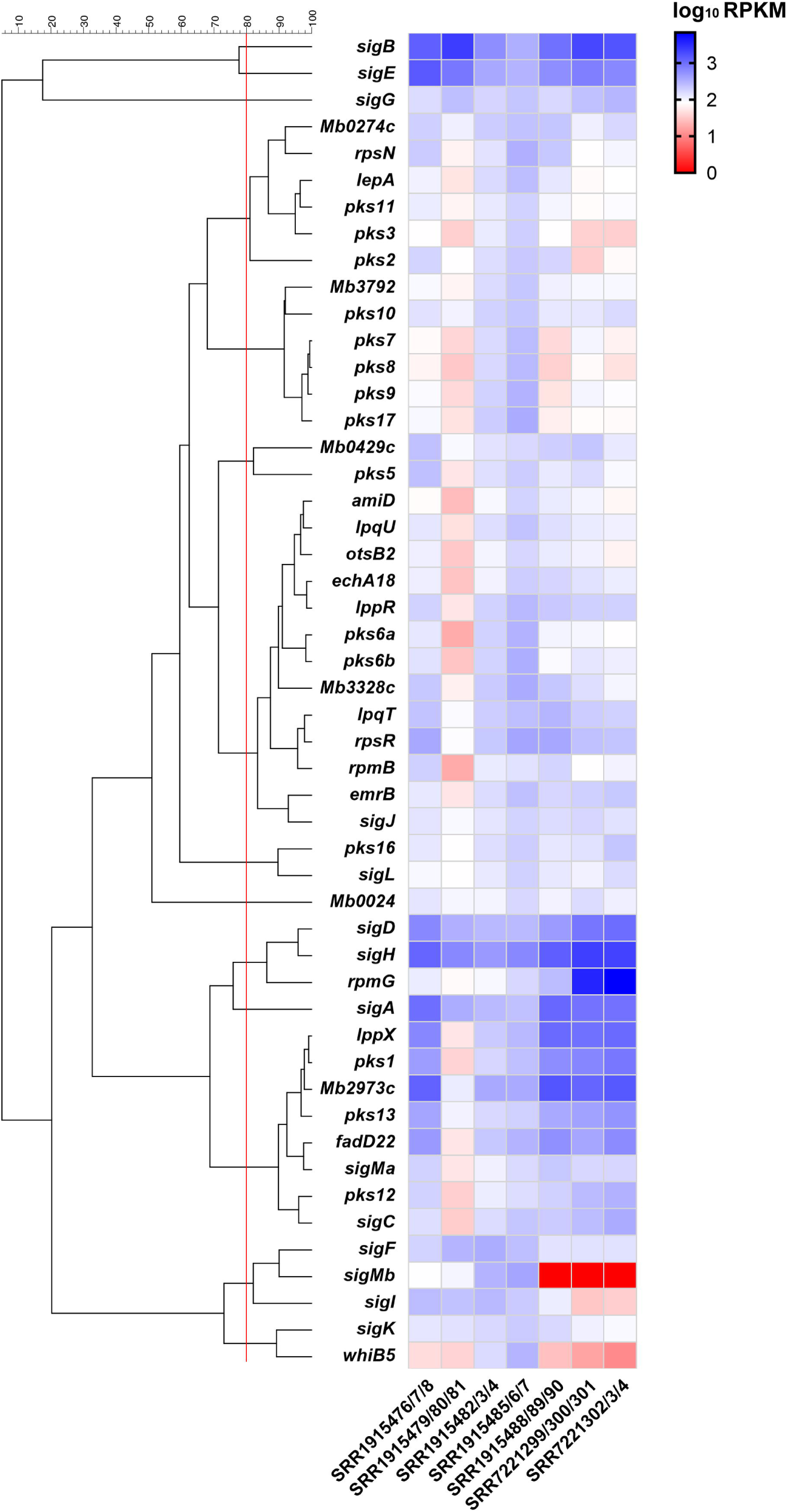
Expression profiling of selected genes from *Mycobacterium bovis*, presented as log_10_ RPKM. Cut-off: 80% of similarity. Abbreviations: SRR1915476/7/8 – *Mb* grown in control conditions; SRR1915479/80/81 – *Mb* grown under starvation for 4 days; SRR1915482/3/4 – *Mb* grown under starvation for 10 days; SRR1915485/6/7 – *Mb* grown under starvation for 20 days; SRR1915488/89/90 – *Mb* after reintroduction of nutrients; SRR7221299/300/301 – *Mb* grown in control conditions; and SRR7221302/3/4 – *Mb* grown with addition of Vitamin B.

### 3.3 Differential expression analyses

As mentioned, pathogenic mycobacteria of the MTC are subjected to a set of different growth conditions while in granuloma and also while exposed to antimicrobial therapy. With this workflow, we were able to validate our analyses, as well as to perform a comparison of the differential expression of the selected genes with regulatory genes, upon previously reported expression. In this analysis, we calculated the differential expression, as log_2_ fold change, of 90 genes amongst which are the genes of interest and 55 genes know to be related to regulatory networks in each of the experimental conditions analysed.

When comparing regular growth conditions with nutrient-depletion and phosphate-depletion (41), the selected set of genes do not present any significant fold-changes in expression. Concerning the regulatory genes, significant fold-changes were only found for *sigB* in phosphate-depletion conditions.

Both for glycerol and pyruvate as carbon sources, *pks1, pks15, fadD22, Rv2949c* and *fadD29* are significantly down-regulated *in vitro* at pH 7, in contrast with the *in vivo* mimicking condition at pH 5.7. In the culture grown in pyruvate, log_2_ fold change values were found to be higher than in the sample grown in glycerol. When comparing carbon sources, there is no significant difference in expression of the selected genes of interest at pH 7. However, at pH 5.7, a significant difference in *lppX* and *fadD29* expression was registered, meaning that those genes are slightly downregulated in glycerol as sole carbon source (Figure 6). As mentioned above, the conditions here explored allow the comparison between basal *in vitro* growth and *in vivo* growth inside phagosomes, wherein pH is lower. It is known that mycobacterial cell wall represents a major barrier to the entry of protons from its surrounds due to its complex structure (42). Also, it is known that most of the acid-sensitive mutants identified in *Mtb* present defects in genes involved in cell wall functions and that several cell wall and lipid biosynthesis are regulated by exposure to low pH (43). Several of the regulatory genes reported for acidic pH response (44) were also found to be induced, namely *pks2, pks3, pks4, papA1* and *papA3* (Figure 6).

**Figure 6.**
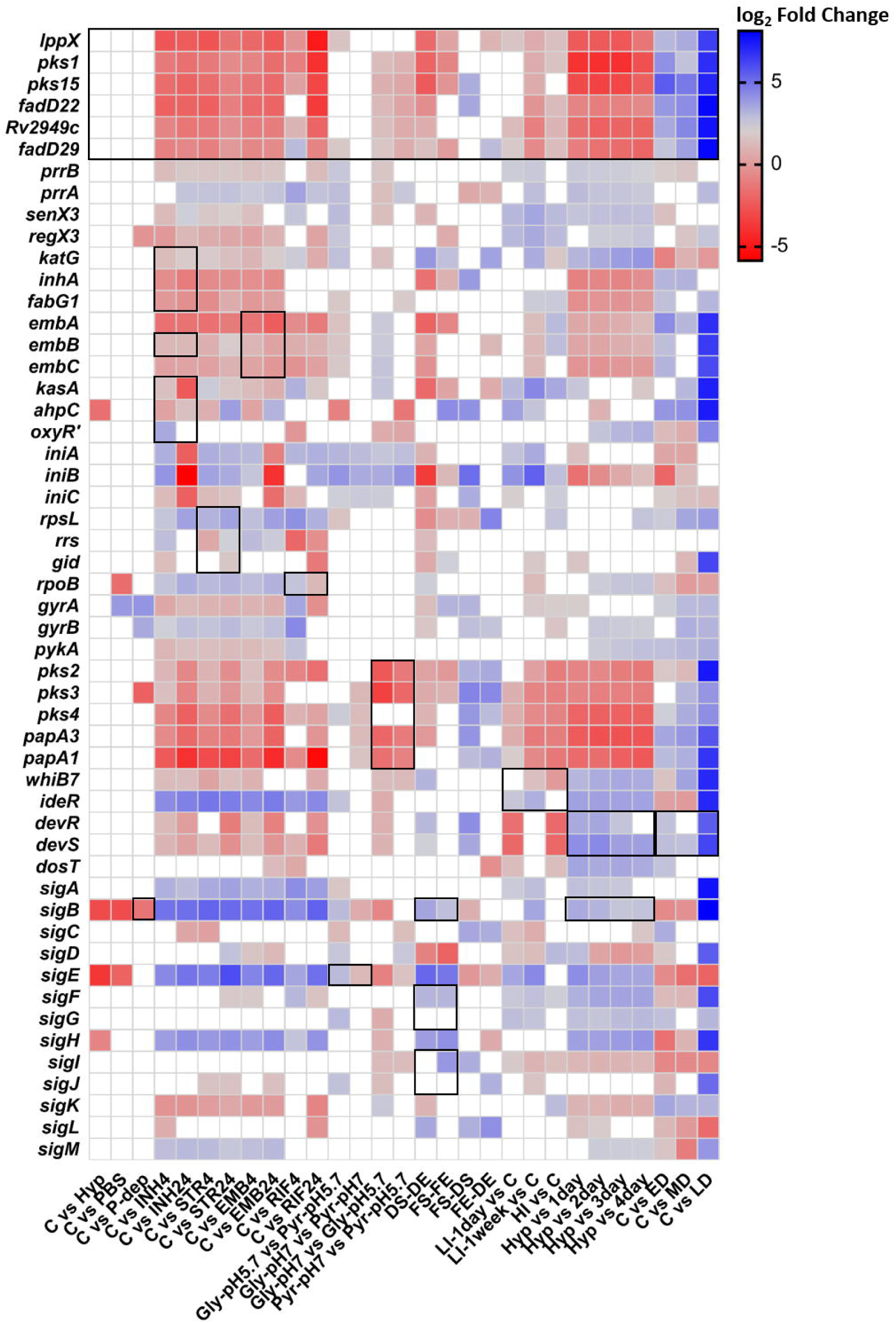
Differential gene expression represented in log_2_ fold change. Most relevant fold changes identified by black outline. Abbreviations: C – Control condition; PBS – *Mtb* H37Rv grown with PBS; P-dep – *Mtb* H37Rv grown in phosphate depletion; INH4 - *Mtb* H37Rv grown with INH for 4h; INH24 - *Mtb* H37Rv grown with INH for 24h; STR4 - *Mtb* H37Rv grown with STR for 4h; STR24 - *Mtb* H37Rv grown with STR for 24h; EMB4 - *Mtb* H37Rv grown with EMB for 4h; EMB24 - *Mtb* H37Rv grown with EMB for 24h; RIF4 - *Mtb* H37Rv grown with RIF for 4h; RIF24 - *Mtb* H37Rv grown with RIF for 24h; Gly-pH 7 - *Mtb* CDC1551 grown in glycerol at pH 7; Gly-pH 5.7 - *Mtb* CDC1551 grown in glycerol at pH 5.7; Pyr-pH 7 - *Mtb* CDC1551 grown in pyruvate at pH 7; Pyr-pH 5.7 - *Mtb* CDC1551 grown in pyruvate at pH 5.7; HI - *Mtb* H37Rv - grown in high iron concentration; LI-1day – *Mtb* H37Rv grown in low iron concentration for 1 day; LI-1week-*Mtb* H37Rv grown in low iron concentration for 1 week; Hyp - *Mtb* H37Rv grown in hypoxia; (1-4) day - *Mtb* H37Rv (1-4) day(s) after reaeration; FS - *Mtb* H37Rv grown in long fatty acids at stationary phase; FE - *Mtb* H37Rv grown in long fatty acids at exponential phase; DS – *Mtb* H37Rv grown in dextrose at stationary phase; DE - *Mtb* H37Rv grown in dextrose at exponential phase; ED - *Mtb* H37Rv in early dormancy phase; MD – *Mtb* H37Rv in medium dormancy phase; LD - *Mtb* H37Rv in late dormancy phase.

The comparisons between growth stages and carbon sources (45) pointed out that, for both carbon sources, *lppX, pks1, pks15* and *fadD29*, are down-regulated in the stationary phase, when compared with the exponential phase. In the cells grown in long chain fatty acids, only *pks1* and *fadD29* display extremely significant down-regulation in the stationary phase, while *lppX* and *pks15* also present significant fold changes (*p* values are shown in supplementary material). By contrast, when bacteria were grown in dextrose, the complete set of genes under analysis was extremely significantly down-regulated in the stationary phase, except for *Rv2949c* and *fadD29* that present lower levels of significance for down-regulation (Figure 6). Since we focused on genes that are part of the biosynthetic pathway of PGL, a significant down-regulation was observed when the cultures enter stationary phase, which was expected and may be explained by the fact that synthesis of the cell wall components is reduced at this time point. Comparing dextrose, the standard carbon source used for *in vitro* growth, with long chain fatty acids mimicking the triacylglycerols available in human cells (45), we could only identify the significant up-regulation of *fadD29*, during the exponential phase and of *pks15* and *fadD22*, in the stationary phase, for the cells grown in long chain fatty acids.

In the iron exposure assays, it was possible to observe that, after 1 day of growth under low iron concentration, only *lppX, Rv2949c* and *fadD29* were significantly differentially expressed, while after 1 week of exposure, the six genes are extremely significantly down-regulated when compared to the culture grown in 0.4% glucose alone. Those results are also supported by the direct comparison between the two cultures exposed to low iron concentration for different periods of time, that show extremely significant up-regulation for the culture exposed for 1 day. Similar to results from low iron concentration, exposure to high iron concentration also showed that *lppX, fadD22, Rv2949c* and *fadD29* are extremely significantly down-regulated and *pks1* is significantly down-regulated (Figure 6). We note that the selected set of genes is highly down-regulated under low iron concentrations, which could be related with the fact that iron takes part in several biological processes inside the cell, being required for cytochromes and other hemoproteins involved in oxygen metabolism. That means that iron deprivation can affect essential cellular processes, inducing a non-replicating state, thus reducing synthesis of cell wall components (46). Although these very interesting results were found, our data analysis does not support the currently described regulation of *ideR* and *whiB7*. Significant differential expression for *ideR* in high iron conditions is not registered, which is in conflict with reports describing its up-regulation (Figure 6) (47, 48). On the other hand, for *whiB7* in low iron conditions, a strong up-regulation after 6 h of exposure is described, a condition for which we do not have comparable data (49, 50). However, after one week of exposure, we found that *whiB7* was significantly down-regulated (Figure 6).

Comparing results obtained when cells were grown in hypoxia with the first 4 days after reaeration (51), it was possible to verify that *lppX, pks1, pks15, fadD22, Rv2949c* and *fadD29* were extremely significantly down-regulated in hypoxia, with some of the higher log_2_ fold change values verified across all assays. Also, *lppX* and *pks1* were found to be down-regulated with extremely significant differences from the first to the third and fourth days (Figure 6). Hypoxia induces many changes in mycobacteria. Both in microaerophilic and anaerobic cultures, *Mtb* is known to develop a thickened cell wall which may be important for adaptation to low oxygen conditions (52). However, the selected set of genes is shown to be extremely down-regulated under hypoxia, agreeing with previously published data (52), and suggesting that maybe the cell wall thickening does not occur in the outer layer. On the contrary, some regulatory genes, members of the DosT regulon, namely *devR, devS* and *dosT* were shown to be highly up-regulated at hypoxic conditions (Figure 6), which is also supported by previous studies (52-54).

In the publicly available experimental data used in our analysis, dormancy was induced by growing *Mtb* in K^+^-deficient medium and, after 14–15 days of culture, adding rifampicin (5 μg/ml) to eliminate dividing bacteria (55). By comparing cells grown to three different states of dormancy with a culture grown to log phase in standard *in vitro* growth conditions, we got the higher fold changes across all assays. That comparison showed extremely significant down-regulation for all genes from the defined set in dormancy conditions. When comparing between states of dormancy, it was possible to see that *pks1* and *pks15* are extremely significantly down-regulated in early dormancy, when compared with medium dormancy. Also, for all genes surveyed, extremely significant fold changes were found between medium and late dormancy (Figure 6). While in dormancy, mycobacteria enter a state of low metabolic activity with alteration of gene regulation in order to accumulate triacylglycerols, loss of acid-fastness and a slower growth rate. These observations explain why the selected genes of interest show a strong down-regulation under dormancy. In agreement with previous reports, and similarly to what happens under hypoxic conditions, *devR, devS* and *dosT*, members of the DosT regulon, were shown to be highly up-regulated in dormancy conditions (Figure 6) (52, 53, 56).

We analysed data from drug-induced stress assays that were performed by growing *Mtb* under exposure to 0.5 μg/ml of INH, 0.5 μg/ml of STR, 1 μg/ml of EMB, and 0.25 μg/ml of RIF, separately (57). For each drug, two time-points, 4h and 24h, were compared in reference to control (no drug). Under exposure to INH, *lppX, pks1, pks15, fadD22, Rv2949c* and *fadD29* are significantly up-regulated at both 4h and 24h. Concerning target genes for INH exposure, it was possible to verify that *inhA, fabG1, kasA* and *ahpC* are up-regulated at both 4h and 24h; on contrary, *oxyR’* is shown to be significantly down-regulated only at 4h of exposure. For *katG* and *embB*, data shows significant but slight up-regulation (Figure 6). Concerning STR exposure, all genes under analysis were found to be up-regulated at both 4h and 24h (*lppX, pks1, pks15, fadD22, Rv2949c* and *fadD29*), by opposition to *rpsL, rrs* and *gid*, which show significant differential expression but with small fold changes (Figure 6). When cells were exposed to EMB, it was verified that *lppX, pks1, pks15, fadD22, Rv2949c* and *fadD29* are significantly up-regulated, at both 4h and 24h. The genes *embA, embB* and *embC* were also significantly up-regulated in the cultures exposed to E*MB*, which agrees with previous published data and validates our findings (Figure 6) (57). The exposure to RIF leads to the up-regulation of our selected panel of genes, with significant fold changes at both 4h and 24h [*lppX* (*p*-value=0.0026; *p*-value=1.5×10^−48^), *pks1* (*p*-value=1.36×10^−35^; *p*-value=4.1×10^−255^), *pks15* (*p*-value=4.49×10^−5^; *p*-value=2.35×10^−79^), *Rv2949c* (*p*-value=0.0261; *p*-value=6.65×10^−46^) and *fadD29* (*p*-value=0.0021; *p*-value=2.27×10^−76^)], with more pronounced fold changes after 24h of exposure (Figure 6).

For *M. bovis* BCG, data from starvation assays were collected after 4, 10 and 20 days and after reintroduction of nutrients (58). For the first three conditions, all genes of interest were up-regulated [*lppX* (p-value=1.5462×10-42; p-value=1.5347×10-17; p-value=3.4984×10-18), *pks15/1* (p-value=3.2164×10-30; p-value=1.3595×10-8; p-value=5.5910×10-8), *Mb2973c* (p-value=3.7632×10-12; p-value=8.8716×10-5; p-value=6.1298×10-10) and *fadD29* (p-value=2.6748×10-5; p-value= 0.0107; p-value=3.3421×10-7)] when comparing to control. The differential expression of genes reported to play a regulatory role under starvation conditions in *M. tuberculosis*, such as *Mb3614c* (p-value=2.7774×10-5), *Mb2076* (p-value= 0.0034), *relA* (p-value= 0.0127), *prrA* (p-value= 0.0125), *senX3* (p-value=5.9579×10-5) and *regX3* (p-value=7.6639×10-12), was also evaluated, whereby after 20 days of starvation up-regulation of these genes was noted (Figure 7). In the assays involving the introduction of vitamin B, no differential expression was evidenced.

**Figure 7.**
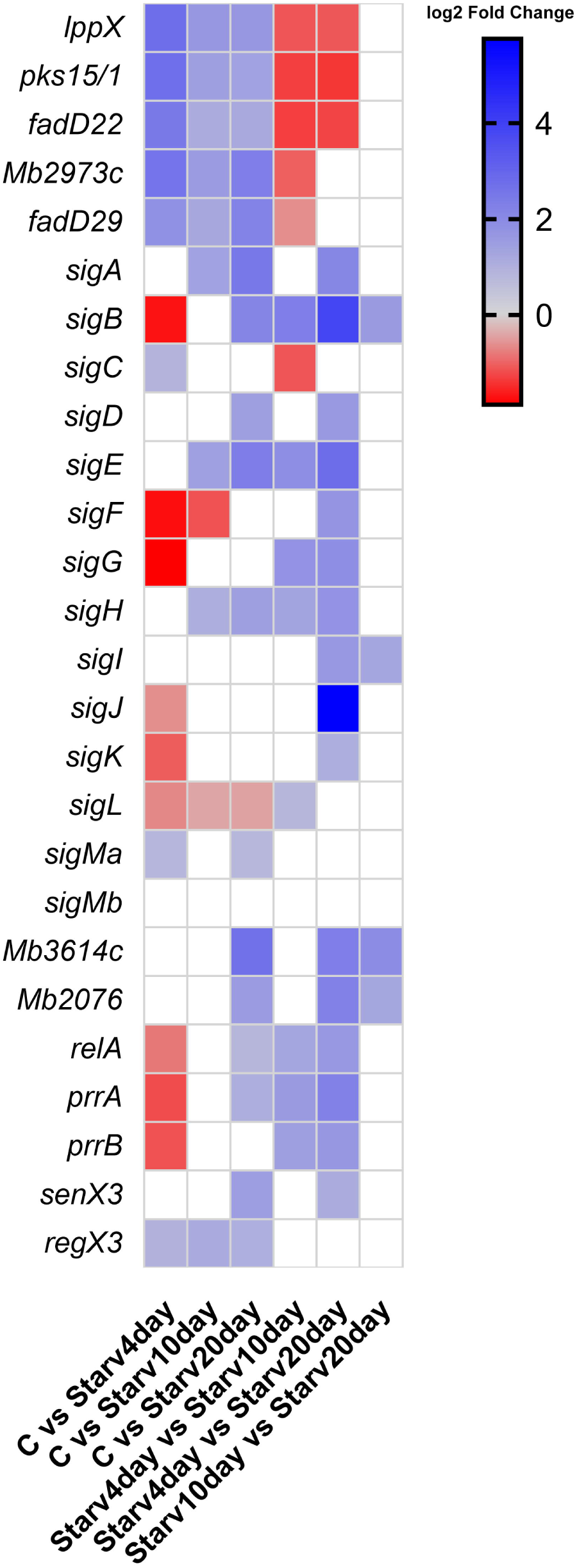
Differential gene expression represented in log_2_ fold change. C – Control condition; Starv4day – *Mb* grown under starvation for 4 days; Starv10day – *Mb* grown under starvation for 10 days; and Starv20day – *Mb* grown under starvation for 20 days.

## 4 Concluding Remarks

*Mtb* virulence is related to its aptitude to survive inside macrophages. During infection, macrophages engulf bacilli, generating a hostile intracellular environment for bacterial replication. *Mtb* interacts with macrophages in a critical and complex process, essential in the arms race between pathogen and host. Recent models of persistency inside the host point to bacterial subpopulations in a latent state that maintain their ability to reactivate upon host’s immunosuppression. One of the processes that is speculated to occur during infection is the development of a cell community structure and organization pattern similar to microbial biofilms. One of the most relevant elements of biofilm formation is the bacterial cell wall, which works as an interface with biotic or abiotic surfaces (15, 59, 60). The study of the regulation of glycolipids biosynthesis such as PGL is of special interest once PGL might be relevant for mycobacterial ability to form biofilms, which may enhance heterogeneity within infecting bacterial subpopulations, enabling resistance and perpetuating infection. PGL production involves several PKS, namely *pks1* and *pks15*, which have been shown to have a critical role in this pathway, since the presence of a mutation that disrupts *pks15*/1 DNA-coding sequence was associated with lack of PGL production in *Mtb* (11). Also, it is known that the reference strain for pathogenic mycobacteria, *Mtb* H37Rv, presents such mutation while other *Mtb* strains circulating across the world maintain this locus integral. Inferring the regulatory pattern of *pks1* and *pks15*, using a genome-wide approach by analysis of RNA-seq data, could does unveil the regulatory patterns controlling phenolphtiocerol and phenolglycolipid production in pathogenic mycobacteria and, indirectly, the downstream processes in which these molecules intervene.

The analysis of expression data gathered from publicly available sources suggested that the target genes selected for this work, *pks1* and *pks15*, may be transcribed as a polycistronic unit composed by three to six genes located both upstream of *pks15* and downstream of *pks1*. All these gene products, except FadD29’, take part in the biosynthetic pathway of the phenolphtiocerol moiety of PGL. Also, *pks1* and *pks15* both seem to be positively regulated by *sigK* and negatively regulated by *sigE*, based on algorithm prediction (24).

By clustering the expression data from more than 100 RNA-seq datasets for *Mtb*, in a robust set of 40 growth conditions, it was possible to correlate *pks1* expression with that of *fadD22, Rv2949c, lppX* and *pks6*, gathering evidence that the first three genes may be part of a polycistronic structure. With a closer analysis, focused on the correlation coefficient values, we were once again able to confirm that all genes thought to belong to the putative polycistronic structure present similar expression profiles. Correlations between these genes were shown to be above 0.85, except for *pks15*. Also, we found that the *pks1* correlation coefficient values were above 0.85 with *pks6* and *pks12*. As noted, *pks15* did not show such high correlation values, although this is mostly due to the presence of several null RPKM values and not to the dissimilarity of the expression profile, since the reads are mapped to *pks1* in strains without the *pks1/15* frameshift (e.g. BCG). In this integrative analysis, it was also possible to relate genes encoding σ factors with the selected genes of interest. While these results must be treated with caution, since σ gene expression may not reflect effective factor activity, we found correlation coefficients above 0.8 between *sigG, sigJ, sigK* and *sigM* and the members of this putative polycistronic structure.

As referred to previously, mycobacteria are subjected to several stress conditions while inside macrophages. By analysing the differential expression of *lppX, pks1, pks15, fadD22, Rv2949c* and *fadD29*, it was possible to define under which conditions these genes are positively or negatively regulated. We analysed expression levels of strains grown under a diverse set of conditions, namely pH, carbon source, growth phase, exposure to limiting or excessive iron concentration, hypoxia, dormancy, phosphate depletion and antibiotic exposure. This analysis revealed that our selected genes of interest are up-regulated at acidic pH and antibiotic exposure and down-regulated at stationary phase, under hypoxia and dormancy, and at both low and high iron concentrations. The combination of two sets of data, i.e. clustering of genes by expression data and differential expression analysis, suggests that *fadD29* may be isolated from the remaining set of genes. Also, in one of the conditions, *fadD29* expression seemed to diverge from that of *lppX, pks1, pks15* and *Rv2949c*, and in another, both *Rv2949c*and *fadD29* expression profiles diverge from *pks1* and *pks15*. Using differential expression analysis, we were also able to confirm that *sigK* shares the expression profile with the selected genes of interest in 88% of the exploited conditions with significant fold-changes; almost the same percentage is also verified for *sigJ*. On the contrary, *sigE* presents approximately 90% of expression profile dissimilarity with the selected panel of genes in the conditions under analysis, with significant fold-changes, as well as *sigB*.

While for *M. tuberculosis*, we managed to gather a robust set of data, for *M. bovis* BCG it was only possible to collect data from seven growth conditions of which three represent regular *in vitro* growth, three represent growth under starvation at three time-points and one represents the addition of vitamin B. This smaller data set led to a lower number of clusters. Interestingly, the members of the putative polycistronic structure cluster in the same group. This analysis is in full agreement with the one performed for *Mtb* as all genes of interest are comprised in the same cluster. Also, expression values of *pks1* appear to be similar in control assays performed in *Mb* and in *Mtb* for which log_10_ RPKM varies between 2.4 and 2.8, indicating that although PGL production may be abolished in *Mtb* strains the *pks1* transcript is similarly expressed in both ecotypes, suggesting a secondary role for this transcript.

Building on the information from previously published reports and the transcriptome data we retrieved, compiled and analysed here, we propose a regulatory model for *pks1* and *pks15*. In this model, we use a conservative approach selecting genes coherently sharing expression patterns and exhibiting functional similarity when considering a polycistronic structure model. For this model, we selected a set of four genes, *pks1, pks15, fadD22* and *Rv2949c*, that fulfil the criteria stated above. Based on differential expression analysis, we also selected a set of three σ factors (σ^D^, and σ^B^ and σ^E^) that seem to be involved in the regulation of *pks1, pks15* and *fadD22* expression (Figure 8). Both σ^K^ and σ^E^ were previously predicted by bioinformatic tools to regulate the genes belonging to this polycistronic structure according to mRNA-based expression levels (61), which is coherent with the hereby reported expression data analyses. However, σ^D^ and σ^B^ exhibit similar expression patterns and thus are also included in our model proposal. The genes encoding factors σ^D^ and σ^K^ were shown to be down-regulated under hypoxia and dormancy, as well as in stationary phase. On the contrary, the genes encoding σ^B^ and σ^E^ factors are up-regulated in the same conditions, being that *sigB* was previously shown to be up-regulated under hypoxia (16). While σ^D^ and σ^K^, that appear to positively regulate the selected genes of interest, belong to the lower level of σ factors regulation, the σ^B^ and σ^E^ factors, which putatively regulate in a negative way the selected genes of interest, belong to a hierarchically upper level of regulation. Also supporting the hypothesis that this putative polycistronic structure is regulated by σ^B^ and σ^D^ are some studies analysing the expression profile of knock-out mutants for those sigma factors. For σ^B^ it was reported by Lee and coworkers (2008) that genes encoding for proteins involved in cell wall processes are highly upregulated in complementation mutants, which is in contrast with our data analyses (62). On the other hand, the Δ*sigD* mutant showed reduced expression of genes involved in synthesis of phospholipids and fatty acids (63). Even though our data analyses suggests that σ^K^ is a regulator of this set of genes, when reviewing the published literature on the Δ*sigK* mutant, no differential expression of our genes of interest was reported (64), leading us to exclude this sigma factor as a hypothetical regulator in our model.

**Figure 8.**
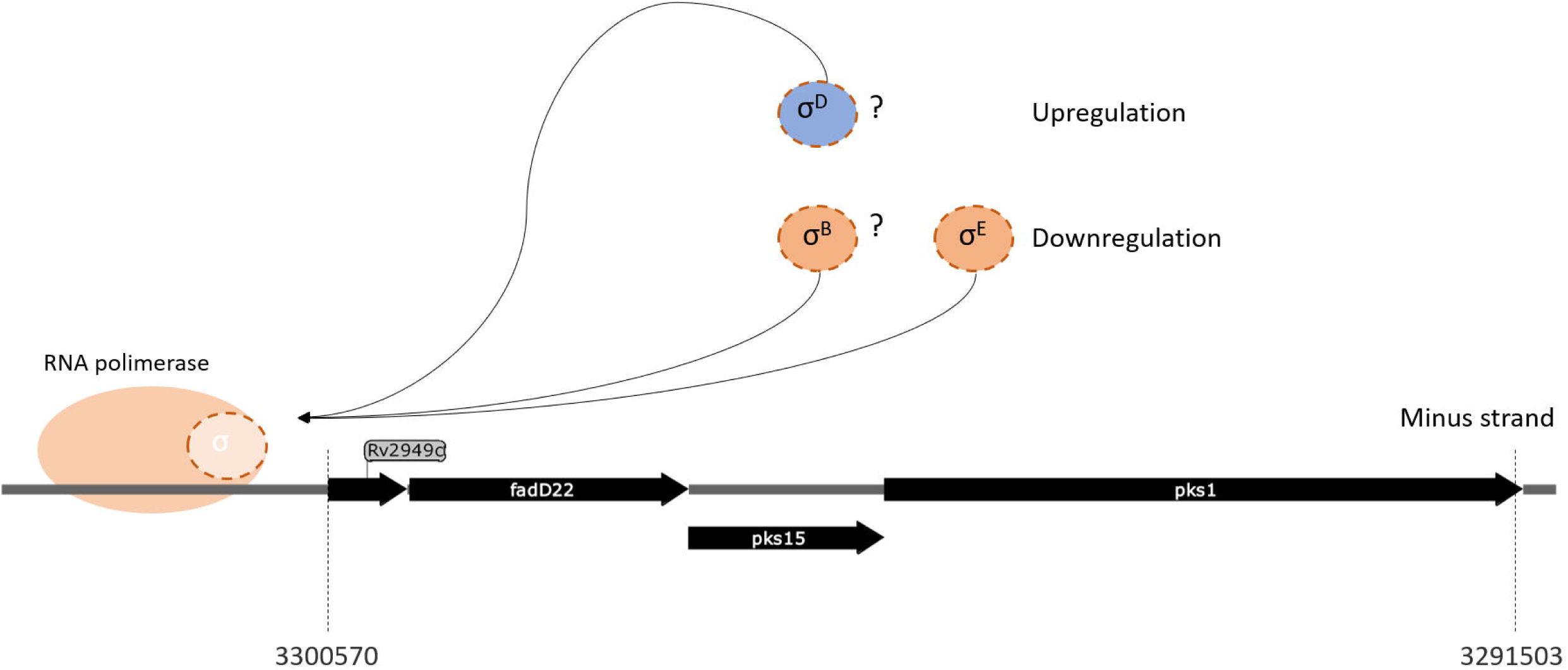
Schematic representation of the proposed polycistronic structure model. The *pks1, pks15* and *fadD22* genes are represented with putative regulation from σ^D^ (positive), and σ^B^ and σ^E^ (negative).

Further experimental validation of the findings described here and of our proposed regulation model may be enabled by classical experiments such as Northern experiments, construction of *pks1* and *pks15* knock-out mutants, and transcriptional fusions with reporter genes of the upstream regions of the genes integrating this putative polycistronic unit in order to unveil the exact location and activity of the promoter. Analysis of *pks1* and *pks15* genes expression in mutant strains of their putative regulators under several growth conditions will further validate the global networks that exert effects on *pks1* and *pks15* activities, previously shown to assume a crucial role in PGL production and thus at the interface with the host.

## 5 Acknowledgements

This work was funded by Programa Operacional de Competitividade e Internacionalização (POCI) (FEDER component), Programa Operacional Regional de Lisboa, and Fundação para a Ciência e a Tecnologia (FCT), Portugal, in the scope of project “Colossus: Control Of tubercuLOsiS at the wildlife/livestock interface uSing innovative natUre-based Solutions” (ref. POCI-01-0145-FEDER-029783), project “MyPATH-Insights into *Mycobacterium tuberculosis* complex PATHogenesis: assessing the benefit of confinement-induced biofilm strategies by comparative genomics” (ref. PTDC/CVT/117794/2010) and strategic funding to cE3c and BioISI Research Units (UID/BIA/00329/2020 and UID/Multi/04046/2020].

## 7 Author Contributions

MVC conceived the study. BR contributed to the conception of methodological approaches and ideas within the manuscript. BR and MVC wrote the first draft of the manuscript. SVG revised and gave critical feedback on all drafts. All authors approved the final version for submission.

## Footnotes

28 Tuberculist available at https://mycobrowser.epfl.ch/; previously available at https://tuberculist.epfl.ch/

29 NCBI available at https://www.ncbi.nlm.nih.gov/

30, 31 TB Database previously available at https://www.tbdb.org/

32 MTB Network Portal available at http://networks.systemsbiology.net/Mtb/

33 PRODIGAL source code available at https://github.com/hyattpd/Prodigal; previously available at http://compbio.ornl.gov/prodigal/ (visited on 11/05/2017)

34 SyntTax available at http://archaea.u-psud.fr/synttax/ (visited on 03/12/2016)

35 NCBI SRA Toolkit source code available at https://www.ncbi.nlm.nih.gov/sra/docs/toolkitsoft/ (visited on 17/05/2017)

36, 37 TopHat source code available at http://ccb.jhu.edu/software/tophat/index.shtml (visited on 17/05/2017)

36, 38 Cufflinks source code available at https://github.com/cole-trapnell-lab/cufflinks (visited on 17/05/2017)

39 ClustVis available at http://biit.cs.ut.ee/clustvis/ (visited on 01/09/2017)

40, 41 Cytoscape available at http://www.cytoscape.org/ (visited on 03/11/2017)

42 htseq-count available at https://htseq.readthedocs.io/en/release_0.9.1/ (visited on 05/08/2017)

46 DESeq2 available at http://bioconductor.org/packages/release/bioc/html/DESeq2.html (visited on 05/08/2017)

49 MicrobesOnline available at http://www.microbesonline.org (visited on 15/11/2017)

